# Brain networks processing temporal information in dynamic facial expressions

**DOI:** 10.1101/819276

**Authors:** Rafal M. Skiba, Patrik Vuilleumier

## Abstract

Perception of emotional expressions in faces relies on the integration of distinct facial features. We used fMRI to examine the role of local and global motion information in facial movements during exposure to novel dynamic face stimuli. We found that synchronous expressions distinctively engaged medial prefrontal areas in the ventral anterior cingulate cortex (vACC), supplementary premotor areas, and bilateral superior frontal gyrus (global temporal-spatial processing). Asynchronous expressions in which one part of the face (e.g., eyes) unfolded before the other (e.g., mouth) activated more the right superior temporal sulcus (STS) and inferior frontal gyrus (local temporal-spatial processing). DCM analysis further showed that processing of asynchronous expression features was associated with a differential information flow, centered on STS, which received direct input from occipital cortex and projected to the amygdala. Moreover, STS and amygdala displayed selective interactions with vACC where the integration of both local and global motion cues (present in synchronous expressions) could take place. These results provide new evidence for a role of both local and global temporal dynamics in emotional expressions, extracted in partly separate brain pathways. Importantly, we show that dynamic expressions with synchronous movement cues may distinctively engage brain areas responsible for motor execution of expressions.

## Introduction

Face perception is a sophisticated skill allowing us to fluently interact with each other (Dols and Russell 2017), recruiting a dedicated and distributed brain network (Duchaine and Yovel 2015). Previous studies using static images of faces have highlighted different perceptual mechanisms that process facial information in terms of both analytic and holistic visual cues characterizing the distinctive spatial configuration of facial parts (Richler and Gauthier 2014; Richler et al. 2008). So called “analytic” visual processes extract information about local facial features, while holistic processes integrate those features into a global configuration. These two mechanisms have been extensively explored by studies of face identity recognition using composite pictures where upper and lower face parts are aligned vs misaligned, or from same vs different individuals (Richler and Gauthier 2014). While processing local parts can subserve face recognition from misaligned or incongruent features in upper and lower parts, an advantage for aligned or congruent features is taken to result from holistic processes sensitive to the whole face configuration.

Similar composite tasks have been used to test emotion recognition with pictures of (static) facial expressions (Meaux and Vuilleumier 2016; Tanaka et al. 2012). These studies suggest that both analytic and holistic/configural processes may contribute to emotion recognition. It is well documented that a smiling mouth region in a face is sufficient to recognize happiness, while the eye region is more crucial for fear or sadness. Other emotions, however, such as disgust or surprise, may require information from both mouth and eye regions (Bombari et al. 2013; Nusseck et al. 2008; Vuilleumier 2005b).

In the human brain, face processing engages distinct cortical areas for the analysis of invariant (e.g., identity) and changeable (e.g., expressions, eye gaze) features, involving ventral visual pathways from Inferior Occipital Gyrus (IOG) to Fusiform Gyrus (FG) and dorsal pathways from IOG to posterior superior temporal sulcus (pSTS), respectively (Haxby et al. 2000). Within this network, holistic processing recruits both IOG and FG (Richler and Gauthier 2014), though the IOG may preferentially respond to local than global face features (Goffaux et al. 2012), unlike the FG (Schiltz et al. 2010). In contrast, analytic processing of local expression features recruits pSTS (Meaux and Vuilleumier 2016; Schobert et al. 2018). In addition, emotional face expressions also activate an extended network beyond these “core” areas, including amygdala, medial and ventral prefrontal areas, as well as sensory and motor areas (Peelen et al. 2010).

Most past studies investigating the role of local and global information in emotion expression used static pictures only focusing on the spatial configuration of face parts. Yet, in real life, expressions dynamically unfold in time, and patterns of coordinated facial movements provide crucial cues for emotion recognition (Fiorentini et al. 2012; Fiorentini and Viviani 2011; Jack et al. 2014). Different emotions are expressed by different combinations of facial action units (FAUs), corresponding to highly specific movements of different groups of facial muscles (Ekman et al. 2002). Accordingly, asynchronous motion among face features hinder emotion categorization performance (Fiorentini et al. 2012), and holistic visual processes may be engaged by dynamic faces to similar degrees as with static faces (Tobin et al. 2016). However, little is known about how motion information is used and integrated across face part to recognize expressions, and which brain pathways are involved. Previous neuroimaging studies of dynamic face expressions have mostly focused on comparisons with static stimuli (Bernstein et al. 2018) or comparisons between different emotions or different social stimuli (e.g., gaze: Schobert et al. 2018). One of the few approaches specifically probing for the role of temporal integration of facial features has employed digitally altered videos in which different face parts exhibit different movement cues with different time-courses, from fully unrelated to fully coherent patterns across parts (Jack et al. 2014; Jack and Schyns 2017; Schyns et al. 2007). In one study combining such stimuli with EEG recording (Schyns et al. (2007), neurophysiology results revealed that the brain integrates local temporal information across dynamically appearing facial parts to form a coherent face percept, with differential responses to motion in the eyes or lip corners starting 50 ms prior to the N170 face-sensitive component. These results also suggested that the integration of emotional expressions may start from the eyes and then move to the mouth, with the N170 peak reflecting the time when the integration of facial parts crucial for categorization of a particular emotion is completed (e.g., corners of mouth becoming characteristic of a happy expression). These data indicate that the brain extracts emotion expression information in a dynamic manner and responds to the relative timing of movement in facial parts that are most discriminative for a given expression.

Here, we used fMRI to investigate neural systems extracting dynamic temporal information from faces using a different approach, derived from the composite face paradigm. Rather than comparing faces whose internal features were spatially aligned or misaligned in a whole face configuration, as with static composite pictures (Meaux and Vuilleumier 2016), we employed digitally-generated stimuli in which dynamic expression cues from the upper and lower face could be either aligned or misaligned in time (i.e., synchronous vs asynchronous). To cover a comprehensive range of expressions and minimize repetitive task demands, our face stimuli displayed four possible emotions (anger, sadness, happiness, joy) that varied in terms of both valence and arousal, as well as in their degree of involvement of the mouth and eye regions. However, in this study, we did not intend to compare the role of different face parts in different emotion categories (Fiorentini et al. 2012; Jack et al. 2014; Schyns et al. 2007). First, we determined behavioral recognition performance and brain activation during the processing of local/asynchronous and global/synchronous expression features in dynamic composite faces, across the four emotion categories. Second, we employed Dynamic Causal Modeling (DCM) to define the functional interplay of areas within the face processing network in response to different expressions dynamics. Our results provide novel insights on neural pathways involved in the processing of motion cues in expressions and their integration across face parts during emotion recognition.

## Methods

### Participants

Twenty-four students of University of Geneva participated in the study (12 females, mean age = 25.9, SD = 5.5). They were paid (30 CHF) for their time in the study. Participants did not have history of neurological and psychotic disease. They reported normal or corrected-to-normal vision. Informed consent was obtained according to local ethics guidelines. All twenty-four participants were included in the fMRI analysis, but nine of them had to be removed from our analysis of eye movements due to poor eye-tracking recording.

### Experimental design

#### Stimuli

Our stimuli consisted of 10 male and 10 female faces generated in FACSGen 2.0 software (Krumhuber et al. 2012) which allows for controlling the magnitude and speed of movement for different facial action units (FAUs) in the upper and lower face parts (Ekman et al., 2002). These facial stimuli expressed four emotion categories (angry, joyful, happy, and sad) that varied along both the valence and arousal dimensions (Feldman, 1995). The four emotional expressions were based on templates provided by FACSGen and only slightly adjusted to balance the duration of videos across conditions.

Our 20 faces were selected from a pilot study using an initial set of 30 identities for which 10 participants (not included in the imaging study) rated emotion expression in the top and bottom parts separately, using an authenticity and an intensity 5-point scales (1- not at all, 5 – very). Only those expressions whose median scores was higher than three on both scales were included in the final set of 20 identities used in the imaging study. An additional behavioral study was also conducted in 46 other observers with a smaller set of highly recognizable expressions (angry and happy) in order to test for recognition latencies and accuracy during emotion categorization across the different types of synchronous and asynchronous stimuli. These data showed no significant differences in emotion recognition performance between the asynchronous and synchronous expressions.

All faces were presented in color, at the screen center, with a resolution of 1200 × 800 pixels. We also used a Control condition where faces had a neutral expression and made one of four possible head movements: left, right, up, and down. Each face expression was presented either with temporally aligned (Synchronous) or temporally misaligned (Asynchronous) motion of expression cues in the upper or lower face parts, as described below. They were displayed using the Psychtoolbox extension (Brainard 1997; Kleiner et al. 2007) for Matlab (The MathWorks, Natick, MA).

In **the Synchronous** condition, the face started with a neutral expression that smoothly transformed (over a period of 800 ms) into one of the four emotion conditions (e.g. angry) with simultaneous changes in upper and lower FAUs, until reaching and maintaining a peak of expression for 2000 ms This was followed by smooth return (over a period of 2000 ms) back to a neutral state (see Figure 1A). The total stimulus duration was 4800 ms.

**Figure 1.**
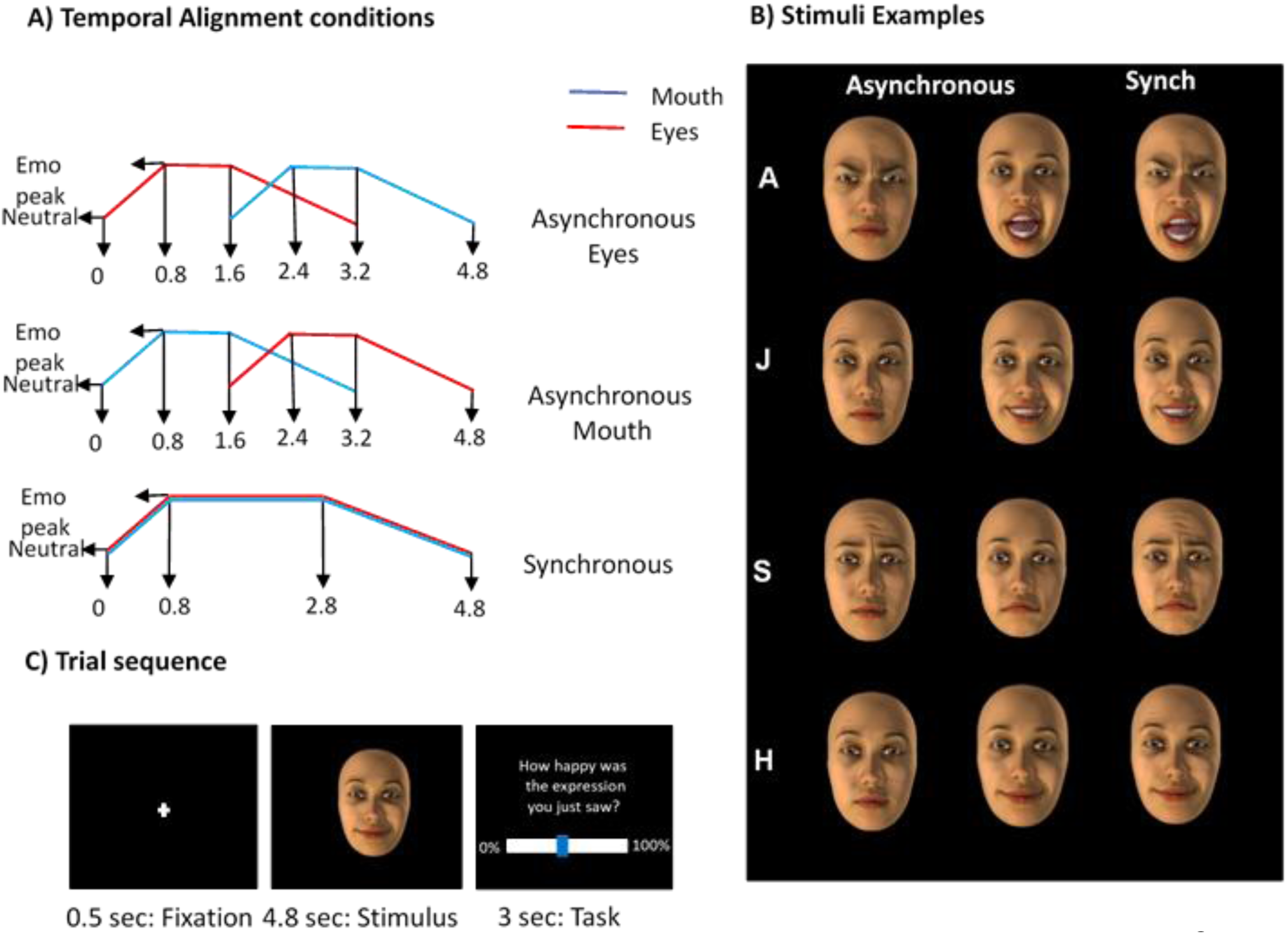
Illustration of temporal alignment conditions, stimuli and trial sequence. **A)** The three diagrams show the temporal unfolding of dynamic facial expressions from neutral to the peak of the expression (Y axis) as a function of time in seconds (X axis), for the three Time Alignment conditions: Asynchronous Eyes (with Eyes first), Asynchronous Mouth (with Mouth first), and Synchronous. Different lines represent different face parts (blue for mouth / lower action units, red for eyes / upper action units). **B)** Examples of the four emotional expressions in each temporal alignment condition. From left to right, the first column shows faces with emotion expression starting in the upper/eye part (here shown at peak intensity while the mouth region is still neutral); the second column shows faces with expression starting in the lower/mouth part (shown at peak intensity while the eye region is still neutral); and the last column shows faces in which both upper and lower parts express emotions synchronously. **C)** Example of a trial sequence (from left to right). Each trial started with a fixation cross at screen center for half of a second, followed by a dynamic face stimulus for 4.8 sec, and a final response display for 3 sec, during which participants had to rate the perceived intensity of emotion expression.

In the **Asynchronous** condition, one part of the face (e.g., eyes) changed first, followed by the other part (e.g., mouth). Half of the trials started with facial movement in the upper part, and the other half started in the lower part. The first movement part unfolded during 800 ms after stimulus onset, remained at the expression peak for 800 ms, and then progressively returned to the neutral state over a period of 1600 ms. The other face part started to express the same emotion 1600 ms after stimulus onset and also reached its peak 800 ms later, then remained at this level for another 800 ms, before returning progressively to the neutral state in 1000ms (Figure 1A). The total duration of these stimuli was 4800 ms, similar to the Synchronous condition.

In the **Control** condition, faces were presented with a neutral expression and a progressive head rotation in four possible directions: up, down, left, or right. For example, in the right condition, the head moved from a front-view facing the participant to a profile view facing the right side of the screen (over a period of 800 ms), then remained in this position for 2000 ms, and finally returned (over 2000 ms) back to the initial front-view position (Figure 1D). These control stimuli also had a total duration of 4800 ms.

Four emotions were displayed in our dynamic expression stimuli: Joy, Angry, Sad, and Happiness. This allowed us to obtain two positive (Happy and Joy) and two negative valenced emotions (Sad and Angry), orthogonally to either high (Angry and Joy) or low arousal (Happy and Sad). Examples of stimuli are presented in the Figure 1B.

#### Procedure

The experimental task comprised 320 experimental trials that represented 4 Emotional conditions × 3 Temporal Alignment conditions × 20 Face stimuli in each, plus 4 Control head movements × 20 Face stimuli each. The trials were divided into five blocks of 64 stimuli, lasting approximately eight minutes. Each trial started with a fixation cross for 500 ms, followed a dynamic face for 4800 ms, and a final rating display of 3000 ms (**Figure 1C**). In the emotional conditions, participants were asked to rate how angry, happy, joyful, or sad the face they just saw was. In the control condition, they rated how “lively” the face they just saw was. Responses were given by using two buttons of a response pad that could move a cursor toward the right or left side of a horizontal scale displayed on the screen (see Fig. 1B). During the experimental task, at the beginning of each block, participants performed a brief 5-point calibration procedure in order to adjust the eye-tracker recording precision.

#### Behavioral measures

Emotion ratings were compared between the four expression categories and the three temporal dynamic conditions. We initially treated the two asynchronous conditions separately (i.e., when expression movements started in the eye vs mouth region) but found no significant differences in ratings (p>.5) and therefore collapsed these two trial types together for all further behavioral analysis. Average rating scores of emotion intensity from each participant were entered as a dependent variable in a repeated-measure ANOVA with two factors: Temporal Alignment (Synchronous, Asynchronous) and Emotion Expression (Angry, Happy, Joy, Sad). Significant main effects and interactions were followed by Bonferroni post hoc t-tests. The Huynh-Feldt correction was applied for conditions that did not meet an equality of variance assumption in the repeated-measure ANOVA here and in following analyses. The data were analyzes using MATLAB (The MathWorks, Natick, MA) and IBM SPSS (IBM Corp. Released 2017. IBM SPSS Statistics for Windows, Version 25.0. Armonk, NY: IBM Corp).

#### Eye tracker and analysis of eye movement data

Gaze position of the participant was monitored throughout the experimental task using an MRI-compatible eye-tracker ASL EyeTrac 6 (Applied Science Laboratories, USA) running with a sampling rate of 60 Hz. To compare conditions, we computed the percentage of time spent with gaze directed to specific areas of interest (AOIs) on the face images (see Figure 3), respectively covering the upper/eye region, lower/mouth region, and nose. Detailed offline analysis of eye movements was conducted on data from 15 participants due to technical issues with the eye-tracker or poor signal in other participants.

As for behavioral data, we first treated all conditions separately with two types of asynchronous expressions. We compared the number of fixation during the first second from stimulus onset, as well as the total time spent in the Eye, Mouth, and Nose Areas of Interest (AOIs). Fixation was defined as an eye position maintained at the same location for at least 100 ms in a box of 5 × 7 pixels. The total time spent in each AOI was defined as a percentage of time spent in this area across a trial, relative to the total stimulus presentation time. These values were submitted to a 4×2×2 repeated-measure (RM) ANOVA using Emotion as a factor with four levels (Angry, Happy, Joy, Sad), Temporal Alignment as another factor with two levels (Synchronous, Asynchronous), and Area of Interest (AOI) with three levels (Eyes, Mouth and Nose). Subsequently, the two Asynchronous conditions (movement starting in the Eye vs Mouth region) were pooled together as we found no significant difference or interaction with Emotion type for these two temporal alignment conditions (all p values > .240). The only dependent variable that showed clear differences between the three Temporal Alignment conditions (Asynchronous Mouth, Asynchronous Eyes, and Synchronous) was the number of fixations during the first second from the stimulus onset. A repeated-measure ANOVA of these data revealed a significant interaction between Temporal Alignment and AOI (F(2.412, 33.768 = 6.710, p = .002, η_p_^2^ = .231), indicating more fixations in the Eye AOI during the first second when expression started in the upper face, and conversely more fixations in the Mouth AOI during the first second when expression started in the lower face.. There was no other significant effects nor interactions (all p values > .073). Therefore, independently of emotion type, when face movements started by either the lower (mouth) or upper part (eyes), they could be reliably tracked by the eye movements of our observers.

#### fMRI measurements

##### Data acquisition

MRI scanning was performed on a 3-Tesla system at the UNIGE Brain and Behaviour Laboratory (Trio TIM, Siemens, Germany). The functional images consisted of 40 consecutive slices parallel to the anterior–posterior commissure plane, covering the whole brain. A T2*-weighted gradient-echo planar imaging sequence was used with the following parameters: repetition time (TR) = 2000 ms; echo time (TE) = 30 ms; flip angle (FA) = 90°; field of view (FOV) = 192 × 192 mm; matrix size = 64 × 64; voxel size = 2 × 2 × 2 mm. The order of slices was ascending. Structural images were acquired with a T1-weighted 3D sequence (MPRAGE, TR/ TI/TE=1900/900/2.27ms, flip angle=9°, PAT factor=2, voxel dimensions: 1 mm isotropic, 256 × 256 × 192 voxels).

##### Preprocessing of fMRI data

Image preprocessing analyses were performed using SPM12 (Welcome Department of Cognitive Neurology, London, UK: http://www.fil.ion.ucl.ac.uk/spm/), implemented in MATLAB version 2015a (The MathWorks, Natick, MA). The first 5 volumes were excluded from analysis to account for T1 saturation effects. The image preprocessing steps included slice-timing correction to account for acquisition time over the whole brain volume, realignment to the mean image of each session by rigid body transformation, co-registration with each participant’s structural image, and spatial normalization to the standard Montreal Neurological Institute (MNI) EPI template, resampled to an isotropic voxel size of 3 mm. Data were spatially smoothed with an isotropic 8mm full-width at half-maximum Gaussian-kernel.

##### Statistical analyses of fMRI data

###### Flexible factorial GLM

A first-level analysis was conducted on data from each participant using the general linear model (GLM) for event-related design in SPM12, with nine experimental event types (4 × Synchronous + 4 × Asynchronous Emotions), plus two other events that controlled for non-specific onset effects of the visual stimuli (Visual condition) and motor responses (Motor condition). An additional analysis included two separate conditions for asynchronous trials starting with upper face movements and those starting with lower face movements, but revealed no reliable difference between these trials (in keeping with behavioral data) and will therefore not be reported in detail. In the individual GLM design, Visual condition was modeled by events with onsets starting at the appearance of the (neutral) face image and one second duration. Experimental conditions were modeled each with their onset starting one second after the stimulus appearance (peak of expression) and a duration of 3.8 sec (until the end of the stimulus presentation). Finally, the Motor condition consisted of two-second long events with onsets locked on the response time. The GLM design also included regressors controlling for head movement-related variance and realignment parameters (x, y, and z translations and pitch, roll, and yaw rotations). Low-frequency signal drifts were filtered using a cutoff period of 128 s. After model estimation, contrast images were calculated for each experimental condition (vs baseline) in each individual participant, and then entered in a second-level group statistics (random effects).

The second level group analysis was conducted on data from our 24 participants using a flexible factorial design with the nine experimental conditions of interest, crossing the factor Temporal Alignment (two levels: Synchronous vs Asynchronous Features) and Emotion Category (five levels: Angry, Happy, Joy, Sad, vs Neutral Control), under the assumption of unequal variance between subjects and between conditions. For all of the reported data we applied FDR cluster correction and p < .001 significance threshold. The corrected FDR cluster size was given by the SPM 12. In some instances when the observed clusters were relatively small we used a Small Volume correction (SVc) of 12 voxels which corresponds to size of Volume of Interests used in our DCM analysis described below.

##### Parametric fMRI analyses

In addition to the analysis above testing for the main effects and interaction of different stimulus conditions, we also conducted two separate GLM models in which we used eye movement data and emotion intensity rating scores as parametric regressors, respectively, allowing us to examine how these factors influenced brain activation to facial expressions.

For the analysis of eye movements, we focused on the Synchronous face condition only because any effect of gaze fixation patterns were not confounded by differences in the visual stimulation itself (unlike the Asynchronous face conditions). In this GLM, at the first level, each emotion trial type (Angry, Happy, Joy, Sad) was combined with a parametric modulator representing the percentage of the total time spent looking at the Eye vs the Mouth region. Higher scores (>50%) on this parameter meant longer duration of fixations to the Eye region in face stimuli, whereas lower scores (<50%) implied longer fixations to the Mouth region. In the second-level group analysis, one sample t-tests were computed to compare the effects of the fixation regressor on brain responses to the different emotional expressions.

For the effect of perceived emotion intensity, both Asynchronous and Synchronous trials were included in a new GLM model with two main conditions (Synchronous and Asynchronous expressions) and parametric regressors based on subjective rating scores given by participants for each trial during the experimental task. On the second level, we again computed one sample t-tests to determine the effects of expression intensity on brain activity and compare these with the effect of stimulus conditions.

##### Dynamic Causal Modeling (DCM)

DCM (Friston et al. 2003) is a method of selecting models of functional brain connectivity based on Bayesian Model Selection (BMS) and allows for direct testing of the strength and direction of interactions between regions of interest (ROIs). DCM is focused on verifying endogenous connectivity (how all nodes in a network are effectively connected), modulatory connectivity (how a specific experimental condition affects connectivity), and exogenous connectivity (how an input connection influences other parts of the network). Here we kept the same input to IOG throughout conditions and focused on testing endogenous and modulatory connectivity in the face processing network as a function of the different stimulus types. We conducted two successive DCM analysis to interrogate both the core face-responsive areas (Kanwisher and Yovel 2006) and the more extended network engaged by emotion processing (Haxby et al. 2000). In both cases, a two-step greedy search procedure was applied (Dima et al. 2011; Ishai 2007; Stephan et al. 2010). The first step used Bayesian Model Selection (BMS) to identify the best (endogenous) connectivity architecture between ROIs, allowing for modulatory effects from all conditions (Synchronous and Asynchronous stimuli) on each node and each connection. The second step examined modulatory connections (by comparing effects of Synchronous, Asynchronous, or both types face stimuli altogether) for those nodes identified in the first step. We used random-effect (RFX) statistics assuming the effects of all conditions are normally distributed in the population.

For the core network, we compared five connectivity models in the first model selection step, followed by a second step with five variants of the best model testing for different modulatory influences by stimulus condition. Each of these models was composed of three core nodes in right hemisphere: IOG (32 −84 12), FG (28 −62 −10), and STS (48 −26 −4). For the extended network, we used the same procedure but now with 13 models to identify the best connectivity architecture in the model selection step, and five subsequent modulatory models to define the effect of stimulus condition on connections. The extended face network was composed of five nodes: IOG, FG, STS, plus AMY (20 −4 −16) and vACC (−2 18 26). The core system cortical regions were selected based on the contrast of All Emotional faces vs Control faces. The volume of interest (VOI) for AMY was defined by our parametric analysis of intensity ratings. The location of the VOI for vACC was based on the contrast of Synchronous versus Asynchronous expressions. We used mostly right-side areas given stronger activations and usual dominance of the right hemisphere in face processing. The exception was the center of the vACC cluster located in the left but spreading to both hemispheres. Each of these regions contained 925 voxel based on a 12 mm sphere that was applied during the VOI extraction. The GLM design used for the both DCM analyses comprised two main condition regressors (Synchronous and Asynchronous expressions), each completed by a parametric regressor (emotion intensity ratings). DCM models were specified using only the two main condition regressors as vectors of interest.

## Results

### Behavioral ratings of emotional intensity

The intensity ratings made on dynamic face expressions across experimental conditions were submitted to a repeated-measure ANOVA with two factors: Temporal Alignment of features (Asynchronous vs Synchronous) and Emotion Category (Anger, Joy, Sadness, Happiness). Importantly, as shown in Figure 2, there was a significant main effect of Temporal Alignment, F(1, 23) = 27.073, p < .001, ηp2= .541. Synchronous expressions were rated higher (average 73.41% of the possible maximum) than Asynchronous expressions (average 57.41%). There was also a main effect of Emotion Category, F(1.587, 36.509) = 26.009, p < .001, ηp2 = .531. Angry expressions were rated higher overall than each of the three other expressions (all p < .001). Happiness was lower than Joy (p < .019) but not different than Sadness (p = 1), while the latter showed only a trend relative to Joy (p = .095). Finally, there was a significant interaction between Temporal Alignment and Emotion Category, F(3, 69) = 7.717, p < .001, ηp2 = .251, which we explored by calculating a difference score between Synchronous and Asynchronous trials for each of the four emotions. An ANOVA on these values not only confirmed a difference between Emotions (F(3, 69) = 7.717, p < .001, ηp2 = 251), but also indicated that the magnitude of the temporal alignment effect was larger for Angry than Happy (p = .011) and Sad expressions (p = .002), but not different from Joy (p = .350). There was no significant differences between the other expressions (all p > .205). Furthermore, a paired comparison between ratings for the Synchronous and Asynchronous revealed significant differences for all emotion categories: Angry (t(23) = 6.127, p < .001), Joy (t(23) = 4.195, p < .001; Sad (t(23) = 3.891, p < .01; and Happy (t(23) = 3.940, p < .01).

**Figure 2.**
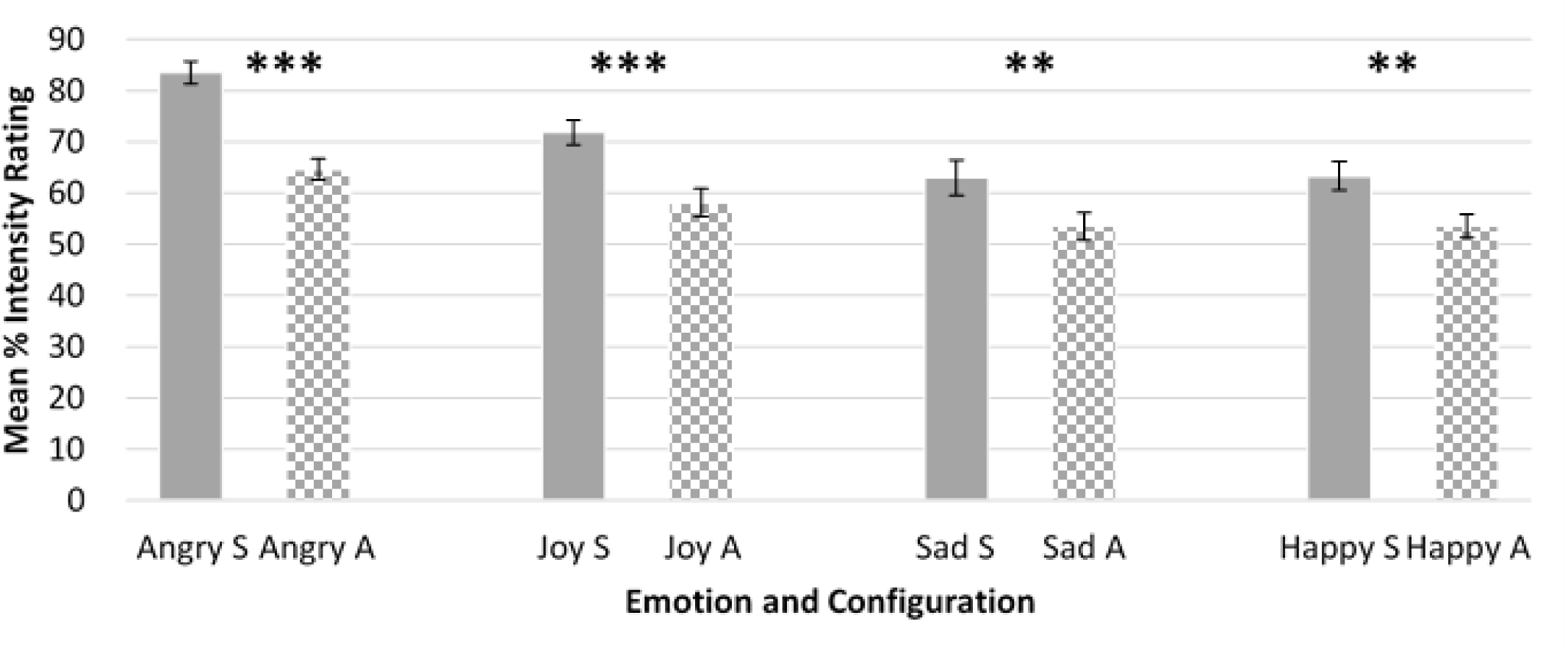
Subjective intensity ratings for dynamic face expressions. Full color bars represent Synchronous trials (temporally aligned movement in upper and lower facial features), whereas checkboard bars represent Asynchronous trials (temporally misaligned condition). Data are shown for each of the four emotions. Asynchronous expressions with movement starting in the upper vs lower face parts did not differ for any emotion category (data not shown). The Y axis shows the mean score of intensity ratings (expressed as percentage on a scale of 100), and error bars show +/− 1 Standard Error of the Mean. ** represents a difference between conditions measured with repeated measure t-test for p < .01 and *** for p < .001.

**Figure 3.**
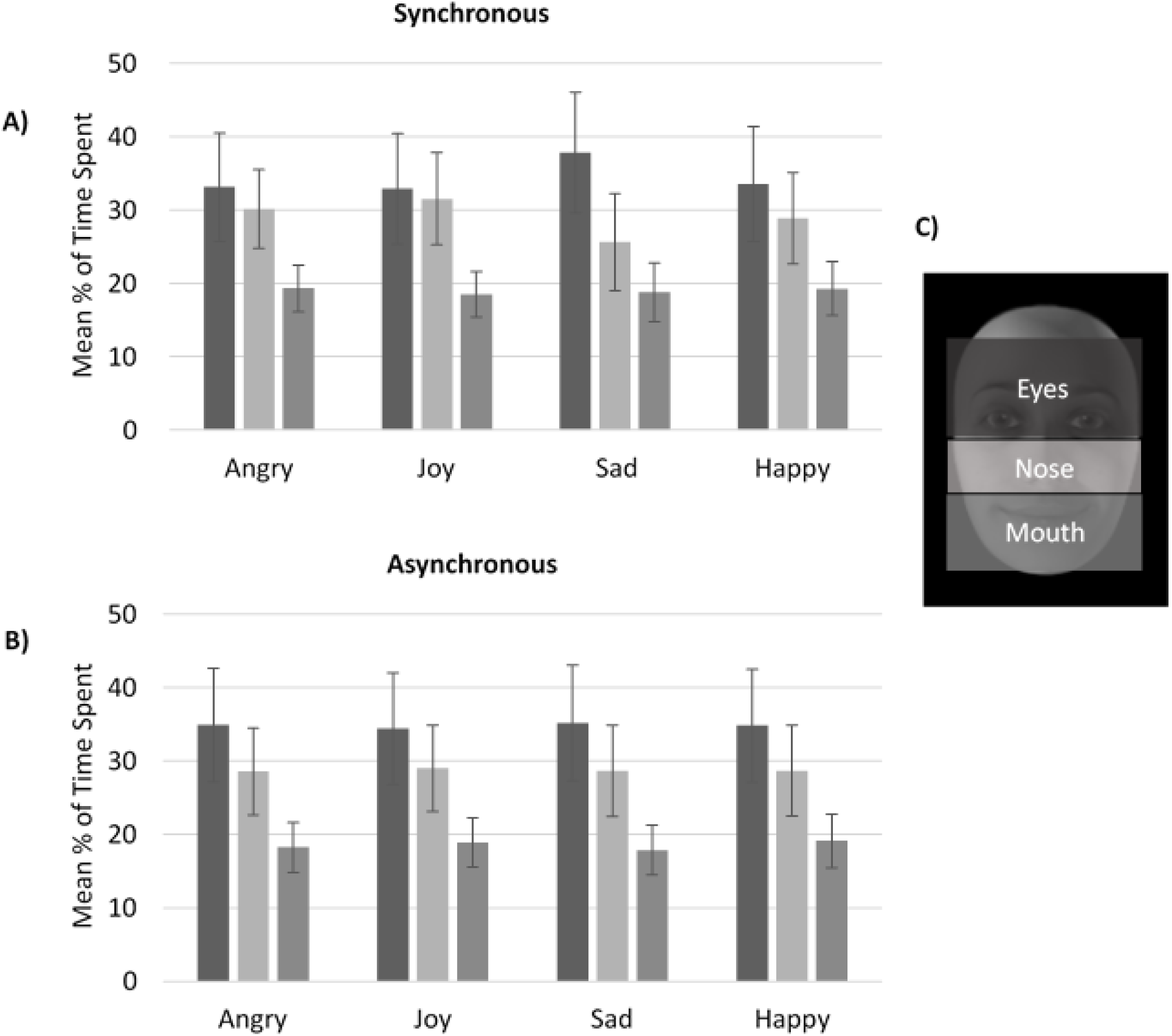
Eye-tracking results. The mean duration of fixations spent in three Areas of Interest (AOI) of face images, during Synchronous (A: top) and Asynchronous (B: bottom) expressions, is plotted for each emotion category. The three corresponding AOIs (upper/eye and lower/mouth parts) are illustrated on a face picture on the right side (C). Additional fixation time was also spent on other image parts, not plotted here. Error bars shows +/− 1 Standard Error of the Mean.

Therefore, overall, Synchronous expressions elicited a perception of more intense emotions in faces across all categories, but this effect was the strongest for the most arousing expressions, particularly Anger, and to a lesser extent Joy (see Figure 2). This advantage of temporally aligned movement among face features thus parallels the holistic effects usually observed for spatially aligned features in static composite faces.

### Eye tracking results

We compared the percentage of time spent on different face parts (upper/eye, central/nose, lower/mouth) across the different stimulus conditions using a 4×2×3 repeated-measure ANOVA (see Methods). This revealed no main effect of any of the three factors (Emotion Category, Temporal Alignment, and Area of Interest [AOI], all F values < 1). There was no interaction between Emotion and Temporal Alignment (F< 1) or between Emotion and AOI (4.933, 64.130) = 1.59, p = .179, η**^*p*^**^2^ = .102). However, there was a three-way interaction between Emotion, AOI, and Temporal Alignment (F(3.164, 44.294) = 2.845, p = .046, η**^*p*^**^2^ = .169). As can be seen in Figure 3, two factors drove this interaction. First, participants looked longer at the eye region than other AOIs in faces with Synchronous Sad expressions (t (14) = .870, p = .399), unlike all other expressions and temporal alignment conditions. Second, they looked at the Eye AOI equally longer than at the Mouth AOI for Synchronous Joy expression, unlike all other conditions. In order to verify those observations we conducted an additional ANOVA on the difference score between Eye and Mouth regions, using the 4 Emotions × 2 Temporal Alignment conditions as factors. This showed no main effect of Emotion (F<1) but the main effect of Temporal Alignment approached significance (F(1, 14) = 3.939, p = .067, η**^*p*^**^2^ = .220). Most importantly, there was a significant interaction between these two factors (F(1.553, 21.748) = 3.917, p = .044, η**^*p*^**^2^ = .219). Further, when inspecting the parameter estimates from this GLM analysis, the largest constant value (i.e., difference from 0) was observed for the Sad Synchronous expressions (B = 12.233), with lower values for other expressions (e.g., the second highest value was observed for the Angry Asynchronous: B = 6.374).

### fMRI results

#### Brain response to dynamic emotional expressions

We first mapped brain areas responsive to emotional face expressions in the main experimental task, by comparing all trials with dynamic expressions (regardless of Temporal Alignment and Emotion Category) against the neutral control condition (head movements). This contrast (Fig. 4) revealed activation in widespread visual regions of occipito-temporal cortex (p < .001, FDRc = 101) including left and right IOG (respectively x, y, z MNI coordinates = −30, −88, 14, Z = 7.54; and 32, −84, 12, Z = 5.67), bilateral lateral fusiform gyrus (28; −62, −6, Z= 4.91; and −28, −68, −14, Z = 4.67), and right STS (48, −26, −4, Z = 5.87). Additional activation was observed in right insula and IFG (54, 20, 2, Z = 5.05), left central motor areas (−34, −23, 62, Z = 4.21), as well as bilateral superior parietal cortex (left: −24, −56, 40, Z = 5.04; and right: 30, −54, 40, Z = 3.98), superior frontal gyrus (left: −34, 46, 24 Z = 4.35; right: 42, 40, 22, Z = 4.16), and right supramarginal gyrus (46, −34, 42, Z = 4.11). Increased activity was also observed in left amygdala but only with a very liberal threshold (Z = 2.19, p < .05 (unc.); −20, −6, −10).

**Figure 4.**
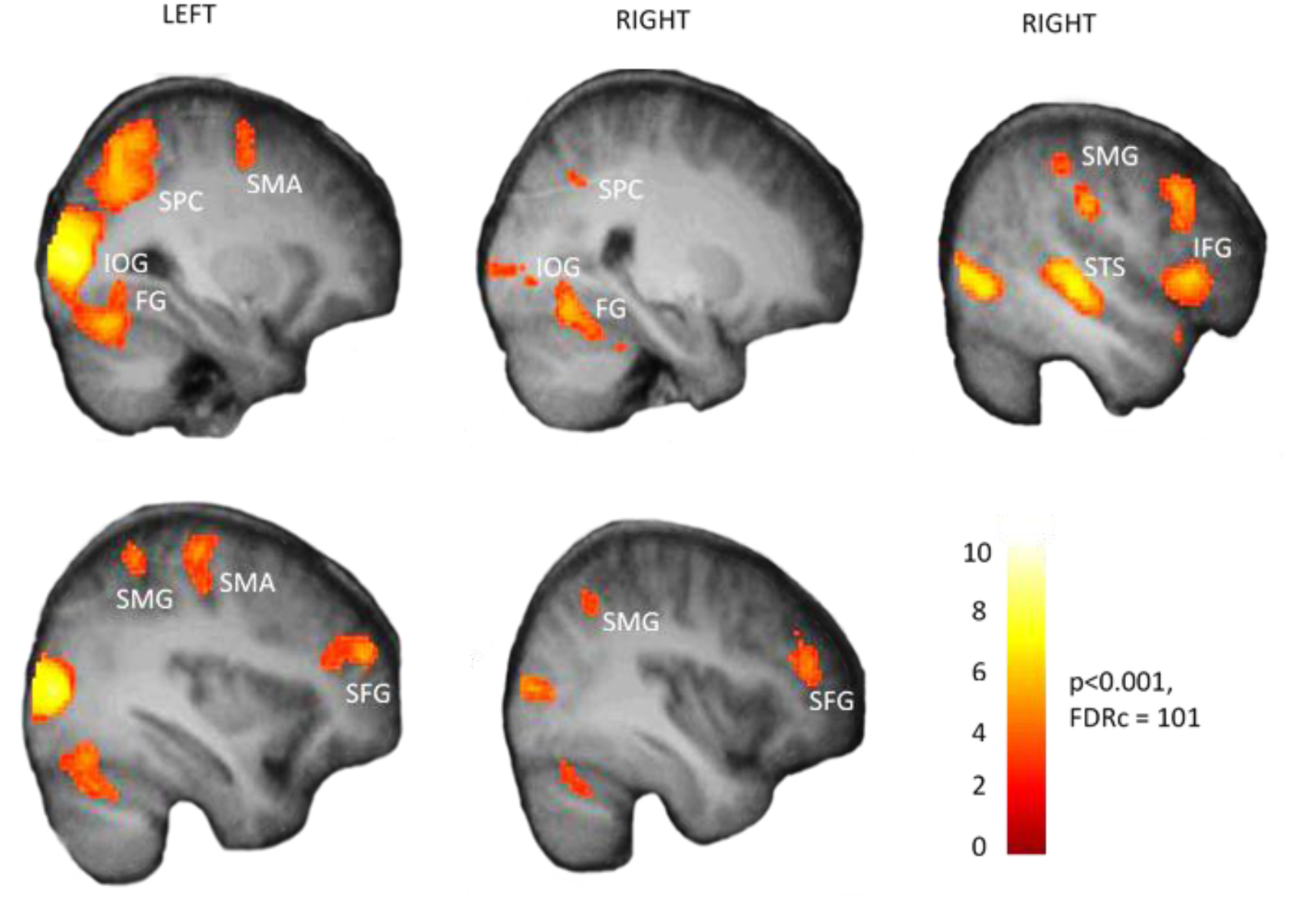
Whole-brain activation maps for the main effect of all emotional faces vs neutral control. Widespread activity is seen in occipital and temporal visual areas, as well as motor and prefrontal regions in both hemispheres. Threshold p < .001 for FDRc = 101 voxels. SPC: Superior Parietal Cortex, IOG: Inferior Occipital Gyrus, SMA: Supplementary Motor Area, SMG: Supramarginal Gyrus, SFG: Superior Frontal Gyrus, STS: Superior Temporal Sulcus, IFG: Inferior Frontal Gyrus, FG: Fusiform Gyrus.

Generally similar brain areas were activated when comparing the Synchronous or Asynchronous expressions separately to the neutral control, in both extrastriate visual areas and prefrontal areas (see **Table 1**). A direct conjunction analysis (set for p < 0.001 FDRc = 116) testing for common increases to Synchronous (vs control) and Asynchronous (vs control) conditions highlighted several regions of the core face processing network (Haxby et al. 2000), mainly bilateral inferior occipital gyrus (left: −30, −84, 6, Z = 7.64; right: 32, −84, 12, Z = 5.17), as well as bilateral posterior fusiform (left: −28, −70, −20, Z = 4.19; right: 28, −62, −6, Z = 4.62), right STS (48, −24, −6, Z = 4.30), and right IFG (58, 20, 0, Z = 4.60), plus left parietal (−24, −60, 44, Z = 4.54) and central-right motor areas (4, 10, 58, Z = 4.33).

**Table 1.** Brain regions activated by Synchronous expressions vs Control (top) and Asynchronous expressions > Control. Coordinates z, y, z are in MNI space. L and R refer to the left and right hemispheres, respectively.

| Contrast | Side | Structure | x | y | z | t | Cluster size |
| --- | --- | --- | --- | --- | --- | --- | --- |
| **Synchronous > Neutral** | L | IOG | -30 | -84 | 6 | 8.299 | 3240 |
|  | L | V5 | -48 | -74 | 4 | 7.751 | ↑ |
|  | R | IOG | 30 | -86 | 12 | 5.531 | 1418 |
|  | R | FG | 22 | -54 | -20 | 5.347 | ↑ |
|  | R | STG | 48 | -24 | -6 | 4.42 | 174 |
|  | L | Pre motor | -36 | -20 | 48 | 5.644 | 1103 |
|  | L | SMA | -8 | 4 | 54 | 5.579 | 1535 |
|  | L | dACC | -2 | 16 | 36 | 4.401 | ↑ |
|  | R | SMA | 4 | 10 | 58 | 4.451 | ↑ |
|  | R | IFG | 54 | 18 | 0 | 4.996 | 376 |
|  | L | SFG | -34 | 48 | 24 | 5.501 | 595 |
|  | R | MPFC | 42 | 42 | 22 | 4.547 | 529 |
|  | R | SFG | 38 | 42 | 14 | 4.162 | ↑ |
| **Asynchronous > Neutral** | L | IOG | -48 | -72 | 4 | 12.646 | 4108 |
|  | L | IOG | -30 | -84 | 6 | 9.001 | ↑ |
|  | L | dPC | -24 | -56 | 40 | 5.385 | ↑ |
|  | R | STG | 48 | -26 | 0 | 7.515 | 1987 |
|  | R | Lateral FG | 48 | -64 | -2 | 7.393 | ↑ |
|  | R | STS | 46 | -36 | 4 | 7.02 | ↑ |
|  | R | PC | 58 | -18 | 22 | 5.479 | 658 |
|  | R | PC | 34 | -54 | 40 | 4.378 | 507 |
|  | R | Pre-SMA | 6 | 14 | 58 | 6.391 | 814 |
|  | L | Pre-SMA | -4 | 8 | 54 | 6.146 | ↑ |
|  | L | Pre-SMA | -28 | -4 | 50 | 4.635 | 223 |
|  | R | IFG | 50 | 18 | 24 | 5.61 | 1620 |

The regions were defined with $p < 0.001$ $FDR_{c} = 101$ . Arrows (↑) means that a region is a part of the same cluster as listed above in the table.

#### Analytic processing of dynamic expressions

We defined brain regions specifically engaged by analytic processing of local dynamic features, regardless of global coherence, by contrasting the Asynchronous expressions to the Synchronous expressions. To ensure that these effects arose in regions sensitive to emotion information, this contrast was computed using an inclusive mask defined by the main comparison of all emotional faces relative to neutral faces (see above).

This analysis revealed selective increases in right STS and right IFG as well as bilateral V5 (Table 2 and Figure 5). These effects of Asynchronous expressions were generally similar in these regions for all four emotion categories when compared with control head movements (see Figure 5B).

**Figure 5.**
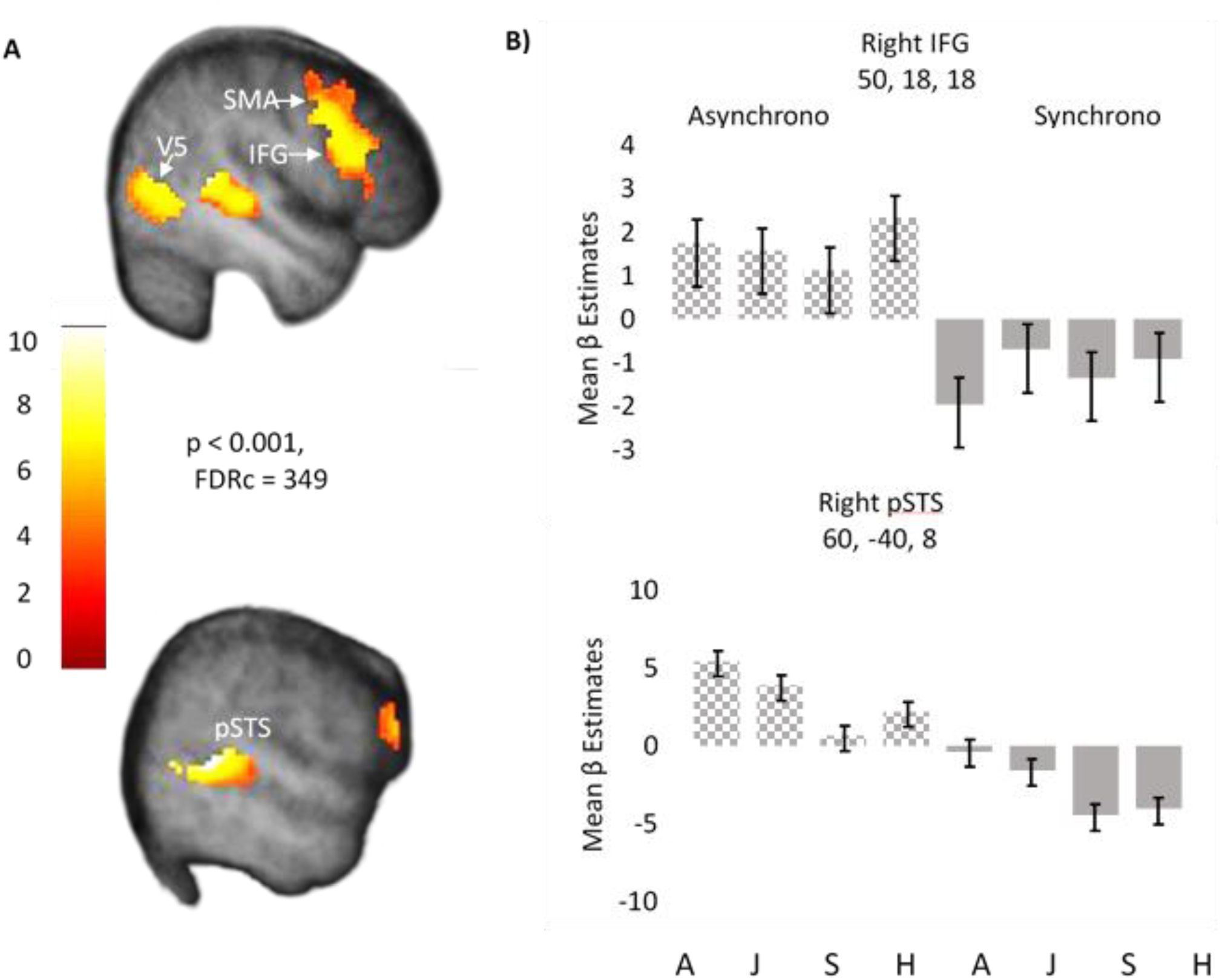
**A)** Whole-brain activation maps showing differential increases to Asynchronous relative to Synchronous expression conditions. Activations (p< .001) were defined within an inclusive mask based on the main effect of Emotion vs Control (p < .05). Significant responses were observed in right STS, V5, and IFG. **B)** Plots showing the mean parameter estimates (beta weights) for responses to each emotional expression (A = Angry, J = Joy, S = sad, H = Happy), extracted from main region identified in this contrast. Error bars show 90% confidence interval of the mean.

**Table 2.**
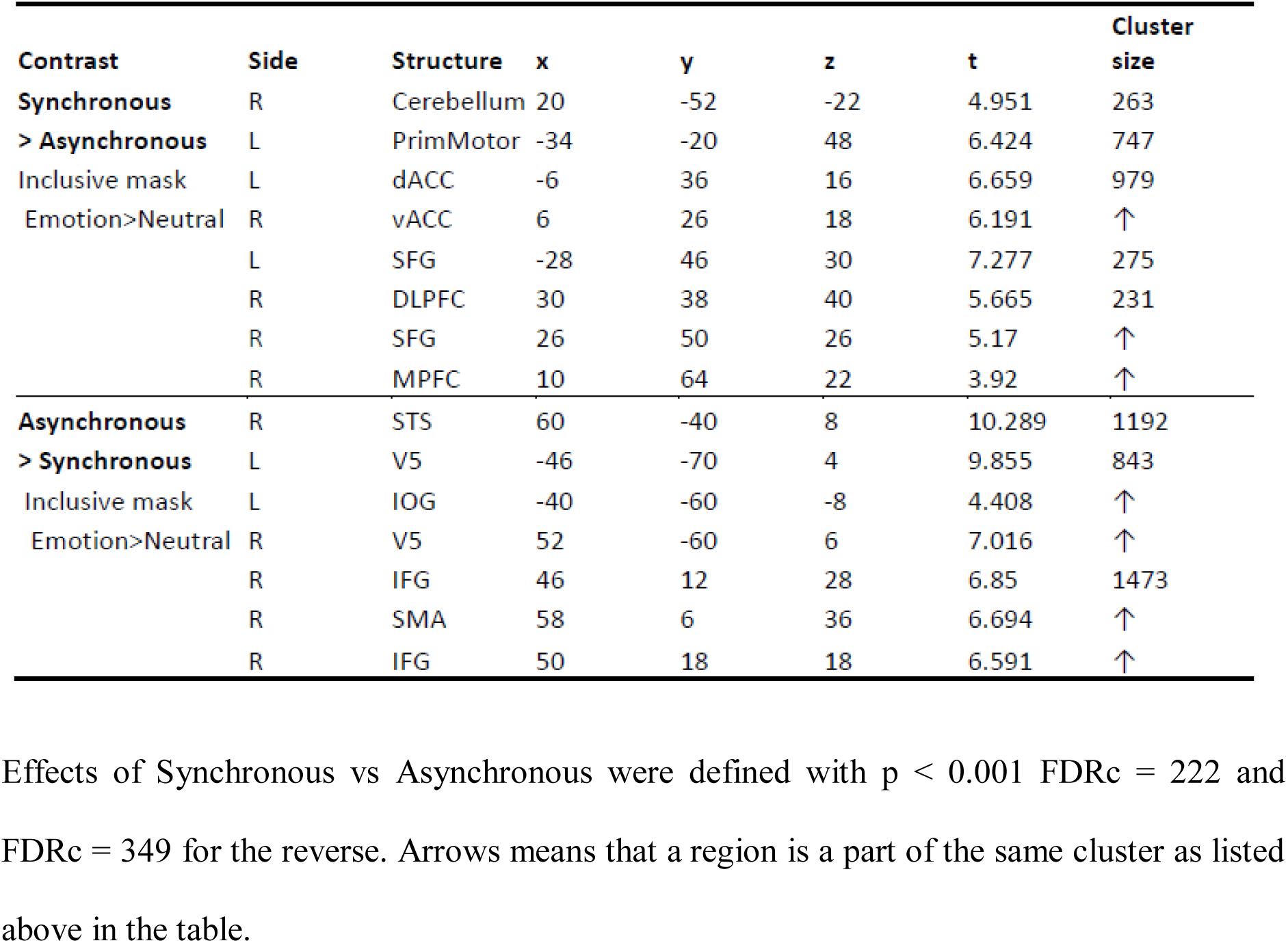
Temporal Alignment effects. Brain areas modulated in contrast of Synchronous vs. Asynchronous (top) and reverse (bottom). Both analysis were masked inclusively by the main effect of Emotion > Control (p < .05).

#### Holistic processing of dynamic expressions

Brain regions specifically engaged by holistic temporal information in dynamic facial expressions were determined by contrasting the Synchronous to the Asynchronous emotional expressions. We applied the same inclusive mask of the main effect of emotion (at p< .05), as described above. This comparison revealed selective increases in several areas within the medial prefrontal cortex, including dorsomedial PFC and dorsal ACC extending to pre-SMA, as well as bilateral regions in superior frontal gyrus (SFG) and central motor cortex (**Figure 6A** and **Table 2**). Remarkably, there was no differential response of visual areas to holistic/coordinated movement among face features. Again, these brain responses were highly similar across all emotion categories (Figure 6B).

**Figure 6.**
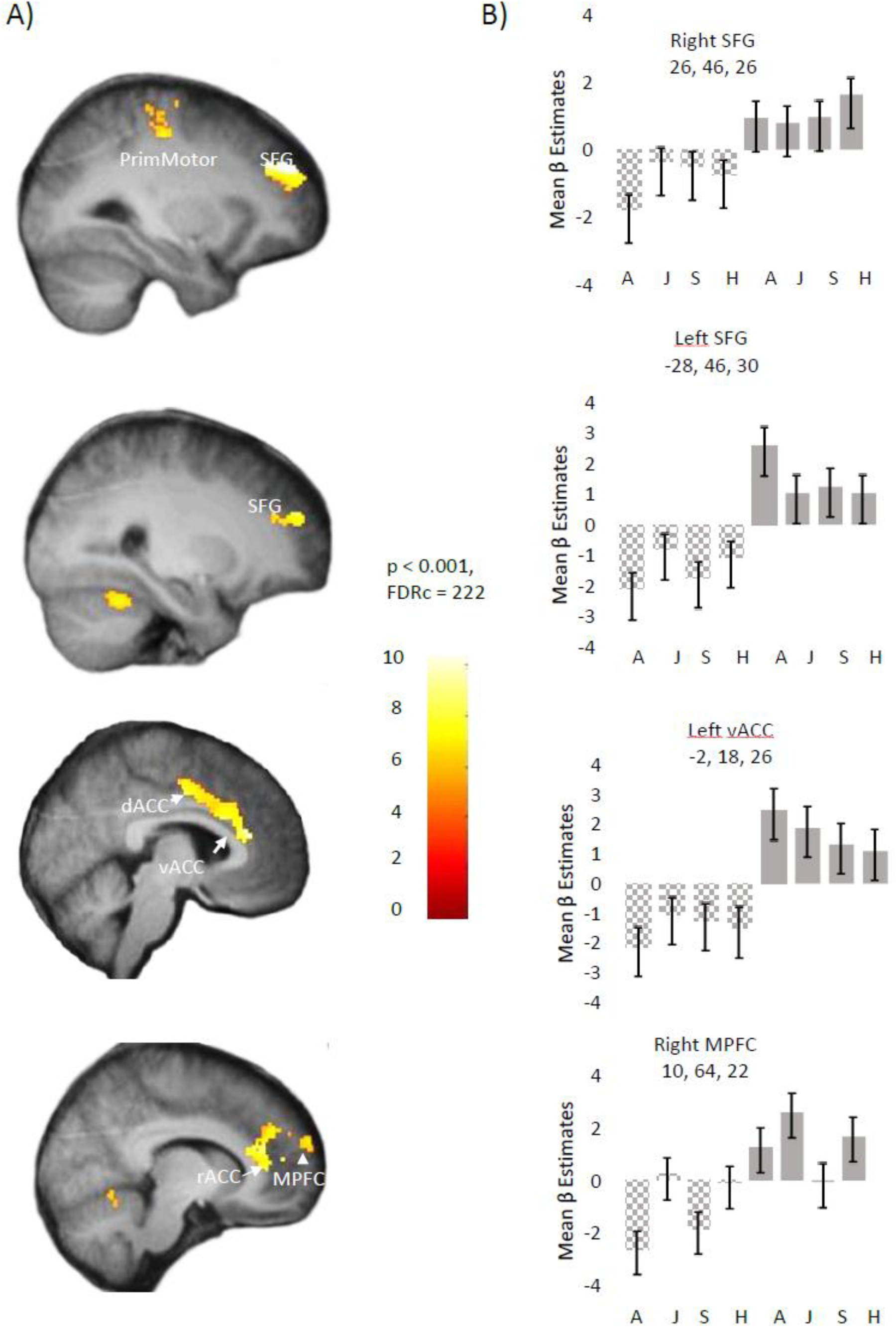
**A)** Whole brain activation maps showing selective responses to Synchronous relative to Asynchronous face expressions. Activations (p < .001, FDRc = 222) were defined after masking inclusively by the main effect of Emotion vs Neutral Control (p < .05). Significant responses were observed in medial prefrontal areas, as well as premotor and lateral motor areas. **B)** Plots showing the mean parameter estimates (beta weights) for responses to each emotional expression (A = Angry, J = Joy, S = sad, H = Happy), extracted from main regions identified in this contrast. Error bars show 90% confidence interval of the mean.

#### Effects of eye movements

To verify whether brain activity patterns were influenced by how participants looked at different parts in dynamic faces, we computed parametric contrasts using the percentage of fixation on to the upper/eye or lower/mouth region as an additional regressor for activations to Synchronous expressions (as there was no confound by difference in movement onset between upper and lower face parts on these trials; see Methods). Results showed that longer fixation times on the Eye region increased activity in right anterior STS (48, 4, −24, Z = 4.46), but far outside the area found to differentially activate during analytic processing of Asynchronous expression (see above). Importantly, there was no effect in other brain areas responding to faces or emotional expressions in previous contrasts. Parametric increases were also found in the bilateral putamen (left: −30, −6, 8, Z = 3.50; 30, −5, 7; Z = 3.53), left caudate (−12, 16, 2; Z = 3.48), and right precuneus (8, −62, 44, Z = 3.52), all areas implicated in eye movement control (Berman et al. 1999). No parametric effect was found for the inverse contrast probing longer fixations on the mouth region. These data indicate that differences observed between Synchronous and Asynchronous expression conditions did not reflect systematic differences in oculomotor behavior.

#### Relation to perceived emotion intensity of dynamic facial expressions

A parametric regression analysis was also performed to assess how activations to Synchronous expressions varied as a function of the perceived emotional intensity reported by participants on these trials. We found higher intensity (across all four emotions) was associated with greater activation of several visual areas in occipital and inferior temporal cortices, including the bilateral IOG and FG, plus right STS. Interestingly, higher intensity ratings were also accompanied by higher activation of the right amygdala and left putamen (**Figure 7**). In contrast, we found no consistent modulations by perceived intensity during the presentation of Asynchronous expressions (except for small increases in V5). This pattern is consistent with the subjectively lower emotional impact of Asynchronous expressions, relative to the more natural Synchronous expressions. Importantly, again, these intensity-related effects did not overlap with differential activations evoked by the Synchronous condition, suggesting that the latter did not merely reflect the higher subjective emotion ratings.

**Figure 7.**
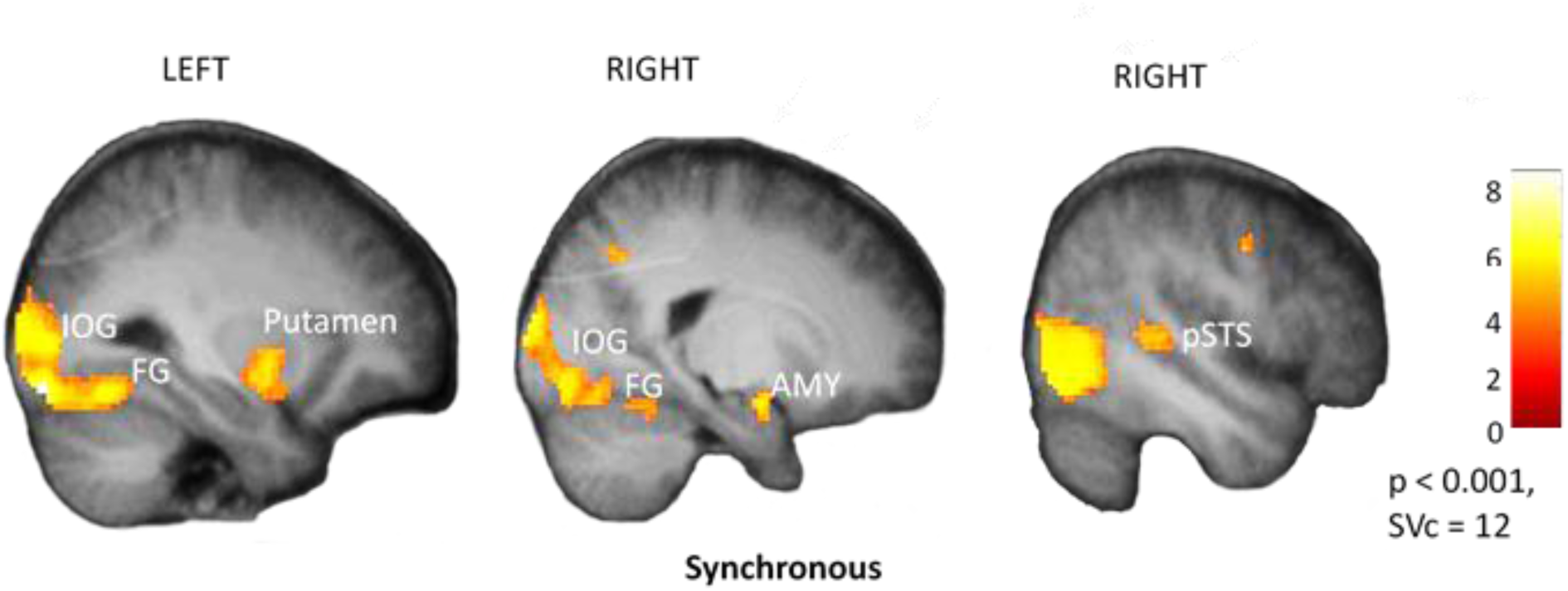
Whole-brain activation maps showing the main effect of perceived emotional intensity from Synchronous face expressions. Small Volume correction (SVc) was applied to face-responsive ROIs, with 12 voxels, p < .001. Significant clusters were present in bilateral IOG (−28, −86, −14, Z= 6.07; and 30, −84, −10, Z = 5.77), bilateral FG (−32, −62, −16, Z= 4.89; and 44, −70, −10 Z = 5.70), right pSTS (44, −34, 4, Z = 4.28), and Amygdala (20, −4, −16; Z = 4.52), in addition to left Putamen (−28, −4, −12, Z = 3.98).

#### Dynamic Causal Modeling (DCM)

In the model selection step of DCM, our connectivity analysis of the core face processing network highlighted two winners among the 5 possible architectures tested (**Figure 8A**): Model #1 and Model #3. Because the latter had a simpler architecture but the same or even higher predictive value than the former, we retained Model #3 for the further analysis. Specifically, we then compared different condition-specific modulations operating on the connections within Model #3 to determine how causal influences between regions varied according to the dynamics of facial expressions (Synchronous vs Asynchronous). As illustrated in **Figure 8B**, the highest exceedance probability was obtained by Model #5 in which information about facial features was transferred in a feedforward fashion from IOG to pSTS, preferentially in conditions of Asynchronous expressions. In contrast, in this model, FG and IOG reciprocally interacted together to exchange information in all conditions equally (Asynchronous and Synchronous).

**Figure 8.**
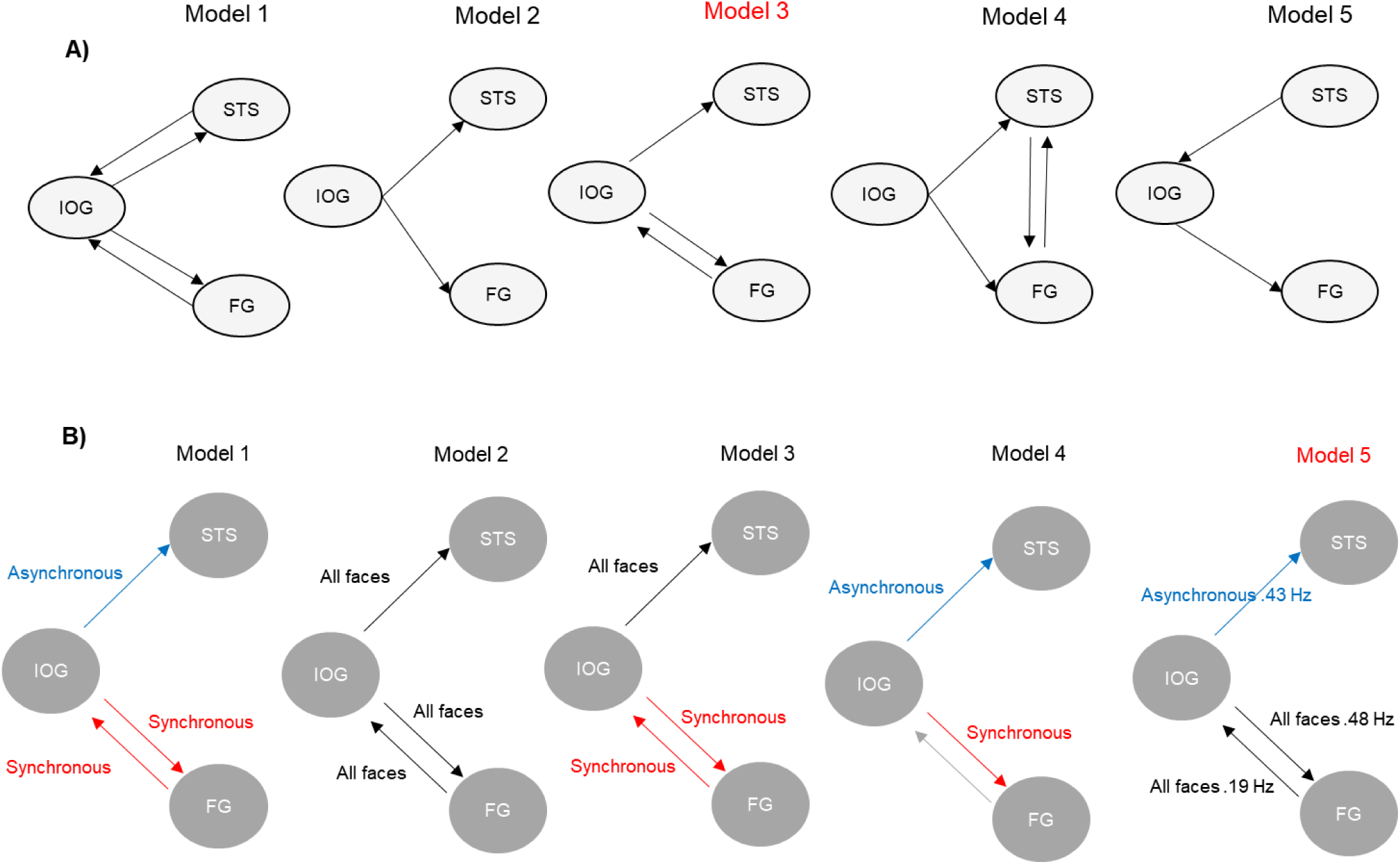
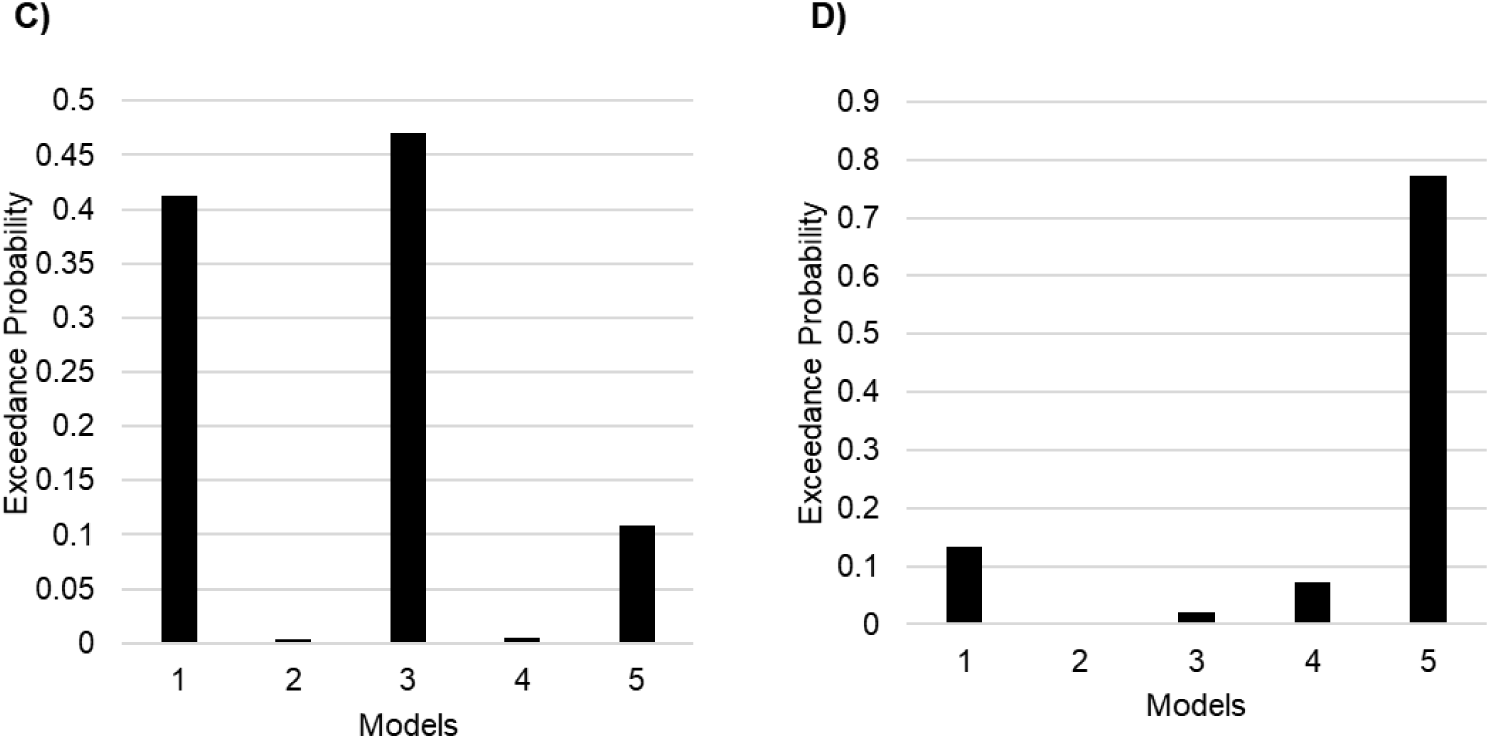
Functional connectivity of the core face processing network in DCM analysis**. A)** Models used to test for the best architecture among core ROIs. Only one input from all face conditions was used to drive IOG, and connections were equally modulated by all four emotion categories. Arrows with one direction mean feedforward connections and double arrows between nodes mean feedback connections. B) Variants of Model#3 from the previous step were used to determine modulations by either Synchronous expressions only (red arrows), Asynchronous expressions only (blue arrows), or both (black arrows). The winning model #5 is depicted with additional measurers of effective connectivity in Hz units. **C)** Bar plots show the highest exceedance probability for Models #1 and #3 among architectures displayed in A. D). Bar plots show the highest exceedance probability for connection modulations in the Model variant #5 among those displayed in B.

Next, we expanded the core connectivity model above to investigate the functional architecture of more extended brain systems responding to dynamic face expressions. We used the same methodology as previously, now incorporating the core regions above in a larger network and testing 13 anatomically plausible models (see **Figure 9A**; only five of these models are shown for simplicity). We compared a variety of connections between emotion-related areas (vACC and AMY) and face processing areas (STS and FG). The wining model was #10, which linked IOG, FG, and pSTS with AMY, as well as IOG with vACC. We then modulated connections in Model #10 according to five new functional models, among which the winner was #4 (**Figure 9B**). These results suggest that only analytic processing of asynchronous expression features was associated with a distinctive information flow, centered on STS, which received from occipital cortex and projected to amygdala. Both STS and amygdala were directly and reciprocally connected to the vACC, where holistic processing of synchronous expression features specifically took place.

**Figure 9.**
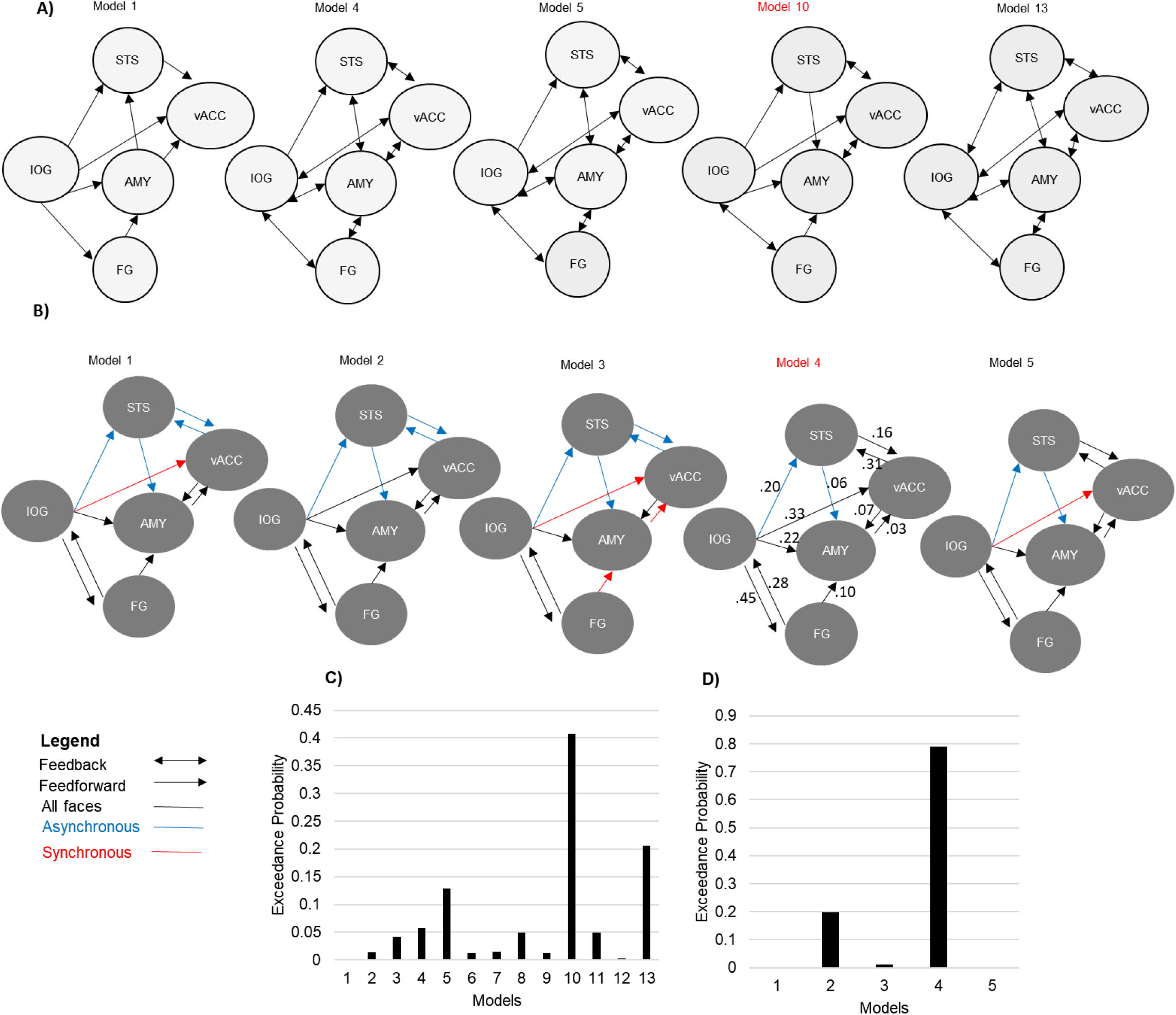
**A)** Connectivity models (out of 13) used to test for the architecture of the extended face network in DCM analysis. Same parameters and conventions as in Figure 8. **B)** Variants of Model #10 from the previous step were used to examine modulations by either Synchronous (red arrows), Asynchronous (blue arrows), or both (black arrows) expression conditions. The wining variant was model #4, depicted with additional measurers of effective connectivity in Hz units. **C)** Bar plot shows the highest exceedance probability for Model #10 among models displayed in A. **D)** Bar plot shows the highest exceedance probability for Model variant #4 among those displayed in B.

## Discussion

This study reveals distinct neural pathways recruited by the processing of global and local temporal-spatial information in facial expressions. Our work extends previous research focused on static relationships with static visual images by using dynamic stimuli and unveiling the role of the temporal alignment of motion cues from different features in coordinated (synchronous) expression movements (as opposed to asynchronous movements). We show that for dynamic facial expressions, emotion recognition processes may not only operate on spatial and metric relationships to integrate facial features into a meaningful “gestalt” (Maurer et al. 2002) but also integrate temporal information across features to extract global motion patterns corresponding to coherent expressions (for a diffrent account of temporal inegration see also: Schyns et al. 2007). This in turn indicates that emotion recognition does not involve only so-called “analytic” processes coding for local dynamics (e.g. in diagnostic face parts), but also engages holistic processes sensitive to global dynamics (across face parts). Accordingly, synchronous expressions were rated as emotionally more intense than asynchronous expressions that included the same local motion cues but misaligned in time, underscoring a perceptual advantage due to global motion information.

At the brain level, this distinction was reflected by activation of different neural pathways. Synchronous processing of dynamic face features (i.e., synchronous expressions) recruited medial prefrontal areas including ventral and dorsal ACC, pre-SMA, anterior DMPFC, as well as motor cortices and bilateral SFG, whereas local dynamic processing without temporal integration (i.e., asynchronous expressions) recruited right pSTS and IFG. Shared activations to both global and local information were found in fusiform and lateral occipital regions, corresponding to the core face network. Further, our DCM analysis revealed that connectivity between these regions was differentially modulated by synchronous vs asynchronous expression cues, with the former projecting from occipital cortex to vACC and the latter projecting to pSTS. These effects were not consequent to higher expression intensity because parametric analyses of emotion ratings showed activations outside prefrontal cortex, specifically in amygdala and visual areas – in accordance with a key role of amygdala in emotional arousal (Anderson and Phelps 2001) and enhanced perceptual processing (Vuilleumier 2005a). Neither were these effects explained by eye movements differences because fixation patterns to upper or lower features modulated only a restricted region in right anterior STS and caudate, outside the expression-responsive areas. Importantly, the eye movements results show that the eye region received more overt attention independently of the presentation condition which accords with Schyns et al. (2007) observation that the integration of temporal face features may start at the eye region and then extend to mouth.

The brain network engaged by the perception of global and local temporal cues in our study are generally consistent with those found in previous work that focused on holistic and analytic processing of spatial feature information in faces for identity (Richler and Gauthier 2014) and expression recognition (Meaux and Vuilleumier 2016) with static photographs. This work suggested that IOG constitutes an entry point for facial information (Richler and Gauthier 2014; Zhen et al. 2013), while FG may provide a hub for holistic processing (Richler and Gauthier 2014; Rossion et al. 2012). Wang et al. (2016) also related IOG activity to local/analytic processing by showing voxels in this region activated to a measure of information segregation, whereas those in FG correlated with a measure of integration. In contrast, both FG and IOG were found to respond to holistic information in static expressions, whereas analytic processing of local (eye/mouth) features distinctively engaged STS and IFG (Meaux and Vuilleumier 2016). Our fMRI data confirm that for dynamic expressions, both local and global movement information is processed in ventral extrastriate areas (FG and IOG), while right pSTS and IFG seem specifically involved in more analytic processing of local and asynchronous visual information. Our DCM results also suggest direct reciprocal interactions between IOG and FG across all stimulus conditions, but preferential engagement of the dorsal IOG-pSTS pathway with asynchronous expressions.

Abundant studies found that pSTS plays a key role in the recognition of changeable aspects of faces, including expressions (Engell and Haxby 2007; Schobert et al. 2018). It also activates to body actions (Herrington et al. 2011) and inferences of others’ intentions (Wyk et al. 2009). Schobert et al. (2018) showed that encoding of dynamic emotion expressions in pSTS may be biased toward internal features (e.g. eyes or mouth), consistent with its sensitivity to featural information in static face expression composites (Meaux and Vuilleumier 2016) and local motion cues in asynchronous dynamic expressions (as found here). With a direct connection to V5 (Bernstein et al. 2018; Bernstein and Yovel 2015) as well as to IFG (Ethofer et al. 2013; Gschwind et al. 2012), pSTS is well positioned to link dynamic social signals with adaptive behavioral responses and actions. IFG often activates during perception of static (Fairhall and Ishai 2007; Ishai 2008) and dynamic facial emotions (Sato et al. 2017), and even vocal emotions (Fruhholz and Grandjean 2013), suggesting a role in higher-level appraisal and monitoring of social interactions (Watanabe 2017).

In contrast, we found that synchronous expression cues specifically modulated a prefrontal network composed of ventral and dorsal ACC, preSMA, DMPFC, and SFG, as well as lateral motor areas. Remarkably, these effects occurred outside visual cortices, but overlapped with parts of the extended face recognition network (Haxby et al. 2000). DCM results further indicated vACC entertains direct and reciprocal interactions with both pSTS and amygdala, presumably allowing this region to integrate information about both local and global expression cues. The ACC is a key component of the limbic system, highly connected with other prefrontal areas, motor regions, as well as down-stream brainstem nuclei (Bush et al. 2000; Watanabe 2017). Midline motor areas in preSMA and ACC are implicated in the generation of non-voluntary facial expressions (Morecraft et al. 2004; Müri 2016; Vrticka et al. 2013) and constitute a direct output pathway mediating automatic mimicry to emotional faces and mirror neuron networks with communicative/social functions (Ferrari et al. 2013; Gothard 2014; Minxha et al. 2017). These regions may also contribute to subjective affect, as shown by electrical stimulation during surgery (Caruana et al. 2018). Further, both ACC and DMPFC are linked to emotion regulation (Bush et al. 2000) and affective theory of mind (Corradi-Dell’Acqua et al. 2013; Peelen et al. 2010), while SFG is frequently activated during emotion recognition (Filkowski et al. 2017) and decision making (Rahnev et al. 2016). Moreover, these regions are anatomically and functionally connected (Watanabe 2017).

Our data thus suggest that decoding of emotion from synchronous facial expressions might partly operate through a recruitment of higher-level motor/premotor mechanisms in medial prefrontal and cingulate cortex (rather than purely visual processes in occipito-temporal areas), pointing to a role of motor and action control systems during holistic recognition of fully coordinated/natural emotion expressions. Such an involvement of motor representations in visual recognition would accord with the notion that perceptual and motor processing are not independent, as shown by psychophysical studies demonstrating a tight interplay between action motor codes and sensory information during stimulus categorization tasks, particularly when motor and sensory processes are coupled through extensive practice (Buckingham et al. 2014). More broadly, this proposal is consistent with cognitive models where visual stimuli are coded along with corresponding actions, such as the ecological theory (Gibson 2014), sensorimotor theory (Noë and O’Regan 2002), or common coding theory (Prinz 1997). A role of motor representations in encoding coherent dynamic expressions may also accord with simulation accounts of emotion recognition (Gallese et al. 2004) and covert mimicry responses when watching facial expressions (Achaibou et al. 2008).

Although providing novel insights in emotion recognition, our study is not without limitations. First, our manipulation of dynamic facial features indirectly probes for holistic temporal processing by using synchrony, but does not directly demonstrate an integrative processes where local perception is changed by global information. Nevertheless, synchronous motion among features increased subjective expression intensity, despite similar visual changes in asynchronous stimuli and similar recognition performance in our pilot/preliminary behavioral testing. Second, by analogy with static composites (Meaux and Vuilleumier 2016), we manipulated only upper and lower facial features but dynamic motion integration might also operate at the local scale among adjacent action units in faces. This more local movement coherence might be processed in different brain areas than the global synchrony examined here. Accordingly, other studies using complex dynamic stimuli with variable combinations of features suggest that temporal integration of expression cues may operate across distinct features within the upper and lower face (Schyns et al. 2007). However, by designing stimuli with asynchronous features across top and bottom parts, we could make sure that participants were exposed to an entire face in all trials while motion information could be extracted from distinct parts in a simple and comparable manner across different emotion expressions, as used in the classic composite task with static pictures (Andrews et al. 2010; Tanaka et al. 2012), and allowing valid fMRI contrast between conditions without visual confounds.

On the other hand, our stimuli might be suboptimal to probe for strictly local processing because all face parts were always presented and could thus engage some aspects of global processing in all conditions. Importantly, our eye-tracking data do suggest that participants differentially focused their gaze on action units relevant for a particular emotion expression over the time-course of a given trial and therefore used analytic processing of local information to some extent. Hence, the current design enables us to probe for relative or preferential engagement of brain networks sensitive to more local or more global motion information in expression while keeping visual inputs generally comparable across conditions.

In any case, the composite paradigm exploited in our study manipulates facial information and motion synchrony in a relatively arbitrary manner across the upper and lower face features. Further research is needed where dynamic expressions are designed with multiple temporal parameters and more fine-grained distribution across face parts in order to identify crucial motion cues underlying the recognition of particular emotions (Jack and Schyns 2017; Schyns et al. 2007). Such stimuli might allow for scrambling temporal aspects of expressions and leave the spatial information intact. However, this approach might have other problems including the generation of unnatural /unrecognizable expressions likely to elicit unclear affective responses in observers and brain activation patterns that would not reflect the normal perceptual processing stream of faces. Furthermore, our main goal here was not to compare different emotion categories or identify specific motion cues associated with particular emotion expressions.

Nonetheless, we found our asynchronous stimuli received lower emotion intensity rating scores. While this supports a perceptual advantage for global synchrony cues, consistent with holistic motion processing, this result may also reflect lower authenticity or lower naturalness of asynchronous expressions that might partly account for increased activity in right STS and right IFG necessary to resolve ambiguity, rather than only mediate local processing of face parts. Please note that the same argument also apply to static composite stimuli that were used in many previous studies and motivated our current design. In addition, it should be underscored that a crucial role of local face processing might precisely be to help resolve ambiguity during face processing, and therefore this system may be needed to support the global system (Bombari et al. 2013; Tanaka et al. 2012). Importantly, as described in our method section, our pilot testing found no accuracy or reaction time differences in the categorization of emotion expressions using the current manipulation of facial dynamics. Therefore, the lower authenticity levels of asynchronous expressions did not make them unnatural in comparison to the synchronous ones.

Finally, we did not record facial EMG during fMRI to determine whether premotor activations to synchronous expressions was associated with involuntary or covert mimicry (Achaibou et al. 2008). However, studies with trial-by-trial EMG measures suggest that spontaneous mimicry correlates with amygdala and orbitofrontal activity rather than medial and cingulate motor regions (Heller et al. 2014). It should be also pointed out that previous research using static images of emotions did not find such strong influence of premotor systems on perception of global expressions (Meaux and Vuilleumier 2016). Therefore, synchronous presentation of dynamic facial action units may affect the motor system and mirror neuron networks in order to create a proper motor plan to express an emotion.

In sum, our novel fMRI results unveil the neural underpinnings of holistic and analytic visual processing of emotional facial expressions in the temporal domain, as manipulated with dynamic (upper vs lower) internal features that were either aligned (synchronous) or misaligned (asynchronous) in time, by analogy with spatial alignment or misalignment in static composites used in previous research (Meaux and Vuilleumier 2016). We show that global/synchronous motion among features engages a medio-prefrontal network bridging limbic pathways with motor areas, suggesting that recognition coherent expression patterns might partly operate through motor codes, in line with a sensorimotor representation of emotion expressions. In contrast, local/asynchronous feature information preferentially engages pSTS and IFG through direct inputs from IOG, whereas the core face processing areas in IOG and FG respond similarly to both global and local dynamic cues. Future research should clarify how these different processes contribute to expression recognition in ecological conditions and for different emotion categories, as well as determine how they are disrupted in brain diseases with affective and social disorders.

## Acknowledgments

We would like to thank Lucas Tamarit for his help with the FACSGen program and Dr. Sven Collette for his help with Matlab scripts. Finally, we are grateful to Prof. Karl Friston for his guidance in the DCM analysis.

